# Improved Parameterization of Protein–DNA Interactions for Molecular Dynamics Simulations of PCNA Diffusion on DNA

**DOI:** 10.1101/2020.02.23.961573

**Authors:** Sunjoo You, Hongeun Lee, Kimoon Kim, Jejoong Yoo

## Abstract

Here, we quantitatively evaluate the accuracy of the protein–DNA interactions in AMBER and CHARMM force fields by comparing experimental and simulated diffusion coefficients of proliferating cell nuclear antigen (PCNA). We find that both force fields underestimate diffusion coefficients by at least an order of magnitude because the interactions between basic amino acids and DNA phosphate groups are too attractive. Then, we propose Lennard-Jones parameters optimized using the experimental osmotic pressure data of model chemicals, by using which one can reproduce the experimental diffusion coefficients. Newly optimized parameters will have a broad impact on general protein–DNA interactions.

## Introduction

Since the discovery of DNA polymerase^1^, numerous enzymes were found to replicate, repair, and transcribe DNA based on the one-dimensional (1D) diffusion along millions to billions of base pairs (bp) long genome^2^. For efficient target searches^3,4^ and accurate gene regulations^5^, living cells control the 1D diffusion of various enzymes such as nucleosome^6^, helicase^7^, single-stranded DNA-binding protein^8,9^ repair proteins^10–14^, and transcription factors^15^. Among those proteins, DNA clamp—the processivity factor for DNA replication^16^, mismatch repair^11^, and chromatin remodeling processes^17^ in eukaryotes^18,19^ and prokaryotes^20,21^—is an outstanding model system to study the diffusion in both experiments^22–24^ and molecular dynamics simulations^19,25,26^.

Proliferating cell nuclear antigen (PCNA)^18,19^—the eukaryotic DNA clamp—forms a homotrimeric complex, in which six protein folds form a closed ring that encircles a doublestranded DNA (dsDNA)^16^. In the inner channel of PCNA, about 30 basic residues—7 lysine (K) and 3 arginine (R) residues per chain—surround negatively charged DNA surfaces (Fig. 1A and 1B). Because the primary role of DNA clamp is providing the processive diffusivity, current models suggest that the K/R residues affect replication and repair processes by controlling the diffusion modes of DNA clamps^27^ Indeed, chemical modification^28^ or mutation^21^ of those K/R residues disrupt the functions of clamps, suggesting that the K/R–DNA interactions are crucial for the function.

**Figure 1.**
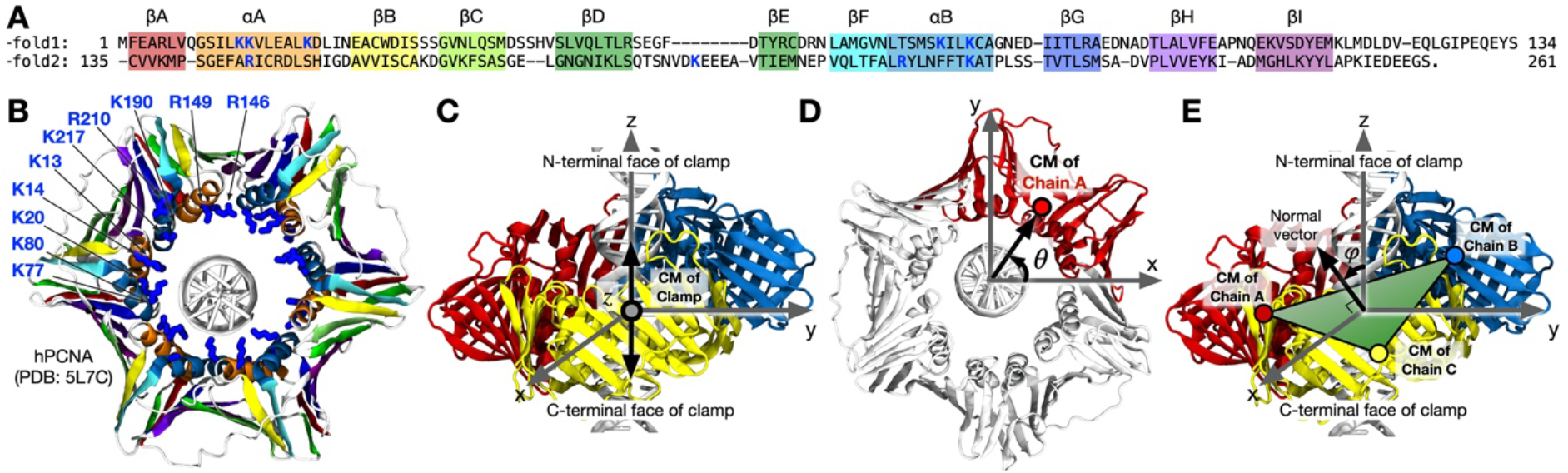
Structure and movements of human proliferating cell nuclear antigen (PCNA). (A,B) The sequence of PCNA monomer (A) and its trimeric clamp structure (B) from Ref.^19^. In panels A and B, each secondary structure and its corresponding sequence are highlighted using the same color scheme. Lysine (K) and arginine (R) residues in the inner channel are shown in blue character in panel A and blue stick representation in panel B. (C–E) Translation (C), rotation (D), and tilt (E) of PCNA are defined by the displacement of the center of mass (CM) of the clamp along the contour of DNA, the azimuthal angle of the CM of chain A projected on the plane normal to the DNA axis, and the angle between the normal vector of the clamp plane formed by CMs of three chains and the DNA axis, respectively.

Here, using PCNA as a model system, we validate the accuracy of K/R–DNA interactions in the molecular dynamics (MD) simulations based on the standard AMBER or CHARMM force field. The test reveals that computed diffusion coefficients of PCNA along DNA are at least an order of magnitude smaller than that measured by the single-molecule experiment^22^. Such unrealistically slow diffusion occurs because the attractive interactions between K/R residues of PCNA and DNA phosphate groups are overly strong for all tested force fields^29^. By optimizing those interaction parameters in the AMBER force field using the experimental osmotic pressure data of model chemicals, we show that MD simulations can quantitatively reproduce the experimental diffusion coefficients.

## Materials and Methods

### Experimental measurement of osmotic pressure

We measured the osmotic pressure of aqueous solutions of ethylguanidinium sulfate, guanidinium acetate, and taurine using Vapro5600 osmometer. The chemicals were purchased from Sigma-Aldrich. For each data point, we performed ten measurements using the auto mode and averaged the ten measured values. The osmometer was calibrated using the standard calibration solutions from Vapro. Entire experimental procedure was performed at room temperature.

### Partial charges of model compounds

We used the standard restrained electrostatic potential (RESP) fitting procedure^30^ to obtain the AMBER-compatible partial charges of ethylguanidinium, sulfate, guanidinium, acetate, and taurine. For each chemical, optimized geometry and electrostatic potential were obtained using MP2/6-31G* and HF/6-31G* in the Gaussian 09 package. The RESP fitting was performed using the AMBER 16 package^31^.

### General simulation protocol

We performed all MD simulations using the Gromacs 2018.2 package^32^. Temperature was kept constant at 300 K using the Nosé-Hoover scheme^33,34^ Pressure was coupled to 1 bar using the Parrinello-Rahman scheme^35^. Van der Waals (vdW) forces were evaluated using a 10-to-12 Å switching scheme. The particle-Mesh Ewald (PME) summation scheme^36^ using a 1.2-Å grid spacing and a 12-Å real-space cutoff was employed to compute long-range electrostatic forces under a periodic boundary condition (PBC). Covalent bonds to hydrogen in non-water and in water molecules were constrained using LINCS^37^ and SETTLE^38^ algorithms, respectively. In all simulations, 2-fs time step was used and coordinates were saved every 20 ps. For all AMBER-based simulations, Joung and Cheatham ion parameters^39^ and the original TIP3P water model^40^ were used. For all CHARMM-based simulations, standard ion parameters and the CHARMM-modified TIP3P water model were used^41^.

### Simulation of PCNA on DNA

We started from the PCNA–DNA cocrystal structure with a 10 bp dsDNA, (ATACGATGGG), that had an exact turn^19^. By duplicating the dsDNA, we prepared a 20 bp dsDNA, (ATACGATGGG)2, that had two exact turns. By aligning the DNA axis to the z-axis and covalently bonding two ends of a strand under the hexagonal PBC (*a = b* ≈ 12 nm, *c* ≈ 6.8 nm, *α = β* = 90°, *γ* = 60°), we simulated an effectively infinite DNA^42^. Protonation states of histidine residues considering the hydrogen bond network in the crystal structure using the pdb2gmx tool in the Gromacs package. The new PCNA–DNA complex was submerged in an explicit 100-mM NaCl solution, energy-minimized for 2,000 steps, and equilibrated for 10 ns with a harmonic restraint (a force constant of 1,000 kJ mol^−2^ nm^−2^) on all non-hydrogen atoms of PCNA and DNA.

### Computational analysis of diffusion coefficients

Because all production simulations were performed with no restraints, every molecule, including DNA, moved freely. Before analysis, we removed the translational and rotational drifts of DNA from the saved trajectory by fitting the instantaneous positions of DNA atoms to their initial positions using the trjconv tool of the Gromacs package. Using the post-processed trajectories, we computed the center of mass (CM) positions of PCNA using the traj tool of the Gromacs package. By processing the CM positions using a custom-made Perl script, we computed translation, rotation, and tilt, described in Fig. 1. It is known that the TIP3P water model overestimates the diffusion coefficients of proteins because the viscosity of TIP3P water is lower than the experimental value^43^. Following Ref.^43^, all MSD and diffusion coefficient data reported in this manuscript were scaled by a factor of 0.375.

## Results and Discussion

### Validation of the standard AMBER and CHARMM force fields

Against the experimental translational diffusion coefficient along the DNA contour of PCNA, *D_t_*, we test the accuracy of the standard molecular dynamics (MD) force fields. Single-molecule measurements of the diffusion coefficient of PCNA showed a mean of 1.16 nm^2^/μs and a standard deviation of 0.79 nm^2^/μs^22^. Given that DNA was stretched to 70% of contour length in experiment^22^, we need to scale the experimental diffusion coefficient by a factor of 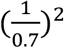 to obtain *D*_t_, resulting in a mean *D*_t_ = 2.4 nm^2^/μs and a standard deviation of 1.6 nm^2^/μs. In MD simulations, we define the translation of PCNA as a displacement of the center of mass of PCNA along the DNA axis (Fig. 1C). Likewise, we define the rotation and tilt of PCNA with respect to the DNA axis (Fig. 1D and 1E).

To simulate PCNA diffusing along linear DNA, we prepared a simulation setup with a PCNA ring encircling a linear 20-bp dsDNA, starting from the PCNA–DNA cocrystal structure^19^ (Fig. 2A). By duplicating the crystallized 10 bp dsDNA that had an exact turn, we created a 20 bp dsDNA that had two exact turns. Then, we aligned the PCNA–dsDNA complex such that dsDNA is parallel to the z-axis and made the dsDNA effectively infinite under periodic boundary condition by covalently bonding two ends of each DNA strand. Because PCNA cannot escape from DNA, this simulation setup is ideal for observing a long time-scale diffusion of PCNA. The simulation box was fully atomistic with an explicit aqueous solution of 100 mM NaCl. See Material and Methods for details.

**Figure 2.**
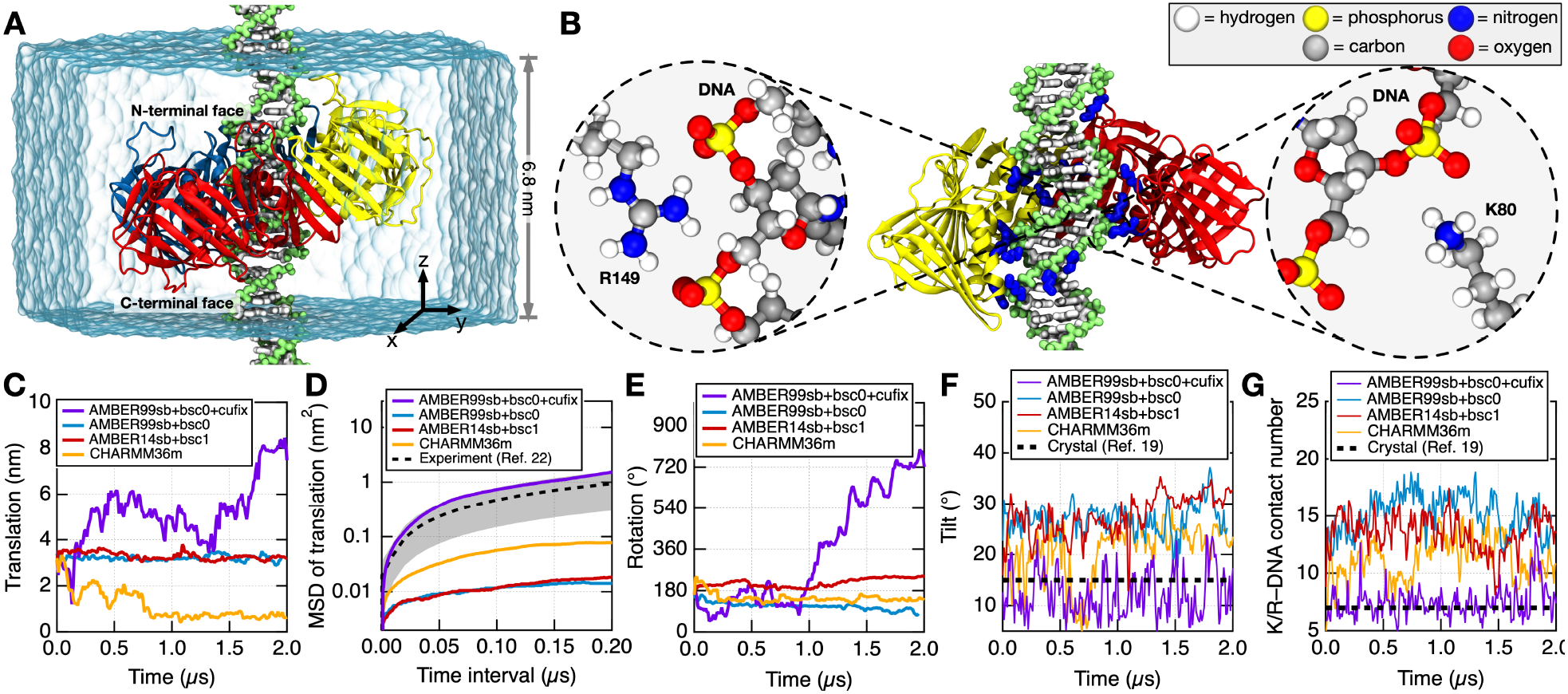
Diffusion of PCNA on DNA in molecular dynamics simulations. (A) Simulation box contained a PCNA complex encircling a 20 base pair DNA^19^, submerged in an explicit 100-mM NaCl solution. By covalently bonding two ends of each strand under the hexagonal periodic boundary condition, we simulated an effectively infinite DNA. Chain A, B, and C of PCNA are shown in red, blue, and yellow cartoon representations, respectively. DNA backbones and bases are shown in green and white molecular representations, respectively. Semi-transparent box indicates the hexagonal water box (ions not shown for clarity). (B) Direct contact pairs between lysine (K) or arginine (R) residues and DNA formed during the simulation using the ff99sb+bsc0 force field. In the center, K/R sidechains in contact with DNA phosphates are shown in blue molecular representations. Chain B of PCNA is not shown for clarity. In the insets, two representative contact pairs are shown in atomistic representations. (C) The translation of PCNA as a function of time. (D) Mean squared displacement (MSD) of translation as a function of time interval, computed using the 2-μs translation data in panel C. Gray dashed line and shade depict the mean and the standard deviation of experimental diffusion coefficient^22^. (E) The rotation of PCNA as a function of time. (F) The tilt of PCNA as a function of time. (G) The number of contacts between K/R residues and DNA as a function of time. A contact is called when a Nζ atom of lysine or a Cζ atom of arginine is within 6 Å from any phosphorus atom of DNA.

Using the simulation setup, we tested three popular force field sets^30^: AMBER ff99sb-ildn-phi for proteins^44,45^ combined with bsc0 for DNA^46^ (hereinafter AMBER99sb+bsc0), AMBER ff14sb for proteins^47^ combined with bsc1 for DNA^48^ (hereinafter AMBER14sb+bsc1), and CHARMM36m. In the 2-μs trajectory produced using ff99sb+bsc0, PCNA stalled at the initial position during the entire simulation, resulting in nearly zero net translation (Fig. 2C and Supplementary Movie 1). To estimate *D*_t_, we computed the mean squared displacement (MSD) of translation as a function of the time interval Δ*t*: MSD = 2*D*_t_Δ*t*. Based on the slope of MSD, we estimate that simulated *D*_t_ is two orders of magnitude smaller than experimental *D*_t_ = 2.4 nm^2^/μs (Fig. 2D). The simulation using ff14sb+bsc1 showed a similar discrepancy (Fig. 2C and 2D and Supplementary Movie 2). In the 2-μs trajectory produced using CHARMM36m, PCNA stalled at a point after a translational drift during the initial 800 ns, resulting in *D*_t_ that is an order of magnitude smaller than experimental *D*_t_ (Fig. 2D and Supplementary Movie 3).

Considering the electrostatic nature of the PCNA–DNA interface (Fig. 1A and 1B), it is highly likely that the discrepancy resulted from overly strong charge–charge attractions between the positively charged sidechains (amine group of lysine, K, and guanidinium group of arginine, R) and negatively charged phosphate groups of DNA^29,49^. Indeed, multiple direct contact pairs between K/R residues and DNA phosphates (Fig. 2B) with lifetimes longer than 100 ns were spontaneously formed for all three force fields (Fig. 3A,B,C). During the simulation, the tilt angle of PCNA increased from 15° observed in the crystal structure^19^ to about 25° (Fig. 2F). The dramatic increase in tilt was a way to maximizing the K/R-DNA contact number as high as 18, significantly deviating from the contact number of 7 observed in the crystal structure^19^ (Fig. 2G). Those artificially stable contact pairs glued the PCNA–DNA interface, stalling PCNA both translationally (Fig. 2C) and rotationally (Fig. 2E).

**Figure 3.**
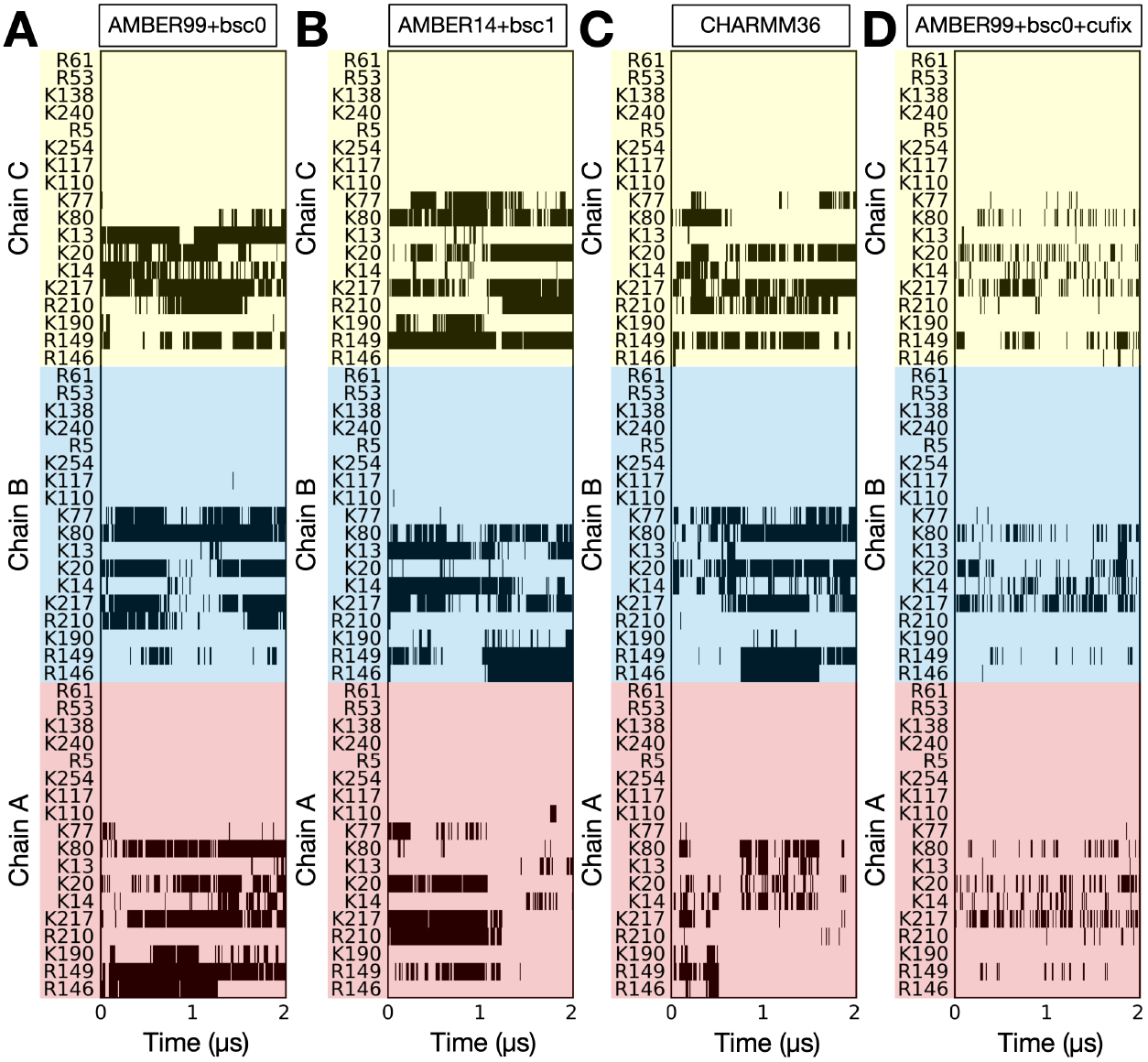
Direct contact pair formations at the PCNA–DNA interface. Residue-specific contact pairs between basic sidechains and DNA phosphate in the simulations using the standard AMBER99sb+bsc0 (A), the standard AMBER14sb+bsc1 (B), the standard CHARMM36m (C), and the AMBER99sb+bsc0+cufix (D) force fields. Black bars indicate that the N_ζ_ atom of lysine or the C_ζ_ atom of arginine is within 6 Å from any phosphorus atom of DNA.

### Calibration of the protein-DNA interaction parameters

Such abnormally strong guanidinium–phosphate and amine–phosphate contact pairs (Fig. 2B) form because of the imbalance between the Coulombic attractions and the steric repulsions described in Lennard-Jones (LJ) potentials^42^. We balanced these pair-wise interactions by calibrating the pair-specific Lennard-Jones (LJ) parameters using the experimental osmotic pressure data as a reference, following the so-called CUFIX procedure^29,49^. Considering that it is generally assumed that sulfate and phosphate are analogs in force field developments, we chose ethylguanidinium sulfate and taurine as model compounds that represent guanidinium–phosphate and amine–phosphate interactions, respectively; see chemical structures in Fig. 4C and 4D.

**Figure 4.**
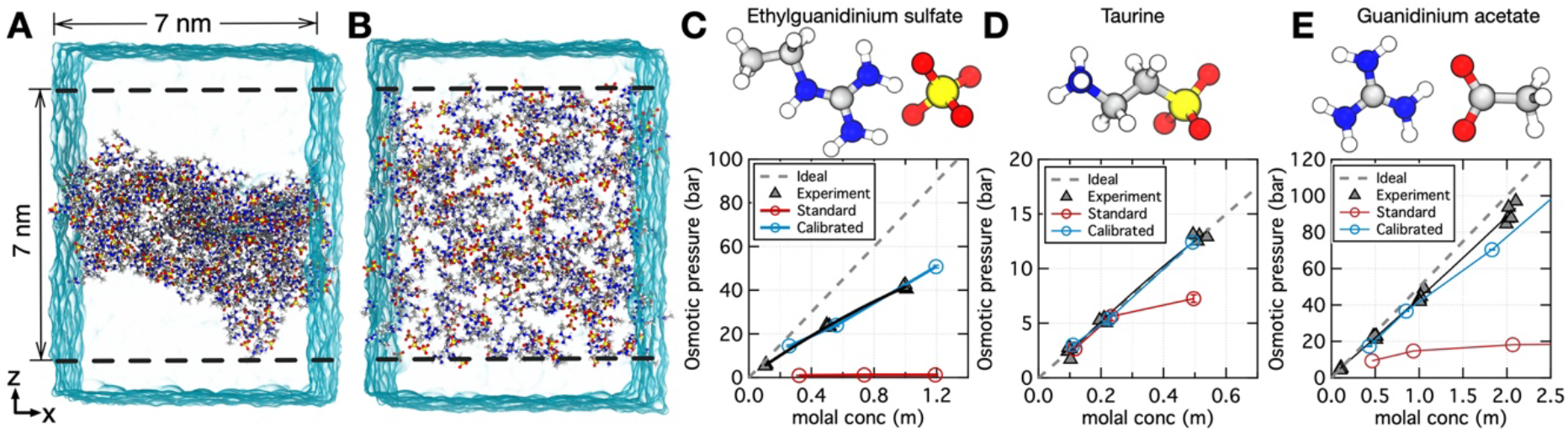
Calibration of AMBER protein–DNA interaction parameters using osmotic pressure data. (A,B) Simulation setup for the measurement of osmotic pressure. A volume of water (blue semi-transparent surface) is divided into two compartments by two planar halfharmonic potentials (depicted by dashed lines) that confine solute molecules within one compartment. Water exchange between the compartments generates osmotic pressure that can be determined from the average force exerted by the solutes on the confining potentials. Panels A and B illustrate instantaneous configurations of a 1.2 m ethylguanidinium sulfate solution observed at the end of 20 ns simulations performed using the AMBER ff99sb force field without (A) and with (B) the calibration. (C–E) Calibration of the Lennard-Jones *σ* parameters for guanidinium–phosphate (C), amine–phosphate (D), and guanidinium–carboxylate (E) interaction pairs using the experimental osmotic pressure of model compounds: ethylguanidinium sulfate, taurine, and guanidinium acetate, respectively. To match the experimentally measured osmotic pressure, we increased the *σ* parameter for nitrogen–oxygen pairs by 0.20, 0.22, and 0.20 Å in panels C–E, respectively. In panels C–E, osmotic pressure data in experiments, MD using the standard AMBER ff99sb force field, and MD using the AMBER ff99sb force field with corrections are shown in gray triangles, red circles, and blue circles. Gray dashed line shows the osmotic pressure of an ideal solution. Note that we took the MD data of guanidinium acetate in panel E from Ref.^49^ to compare it with the experimental osmotic pressure obtained in this manuscript.

We computed the osmotic pressure of the solution of model compounds using a two-compartment simulation setup, Fig. 4A and 4B, in which two virtual semi-permeable membranes (dashed lines in Fig. 4A and 4B) divide a rectangular simulation volume into two compartments. We realized the semi-permeable membranes by using two planar half-harmonic potentials that exert forces on non-hydrogen atoms of the solutes. During the simulations, we recorded the instantaneous forces applied to the solutes by the half-harmonic potentials. Then, the osmotic pressure exerted by the solutes on the semi-permeable membranes was evaluated using the recorded force values. The reader interested in the detailed simulation protocol is referred to the recent review in Ref.^47^.

When simulated using the standard AMBER99sb force field, ethylguanidinium sulfate aggregated into a single large cluster at all concentrations of 0.3, 0.7, and 1.2 m; see Fig. 4A for the representative aggregate at 1.2 m. Consequently, computed osmotic pressure values were nearly zero at all concentrations, contrary to the significantly high experimental osmotic pressure (e.g., 50 bar at 1.2 m), Fig. 4C. To balance the overestimated electrostatic attraction, we increased the steric repulsion by increasing the LJ *σ* parameter for the guanidinium nitrogen and sulfate (or phosphate) oxygen atom pair. As the *σ* parameter increased, the computed osmotic pressure gradually increased. When the *σ* parameter was increased by 0.20 Å, the computed and experimental osmotic pressure values matched at all concentrations (Fig. 4C). Similarly, we increased the LJ *σ* parameters for the amine nitrogen–sulfate/phosphate oxygen atom pair by 0.22 Å to match the experimental osmotic pressure data of taurine (Fig. 4D).

We combined these newly developed corrections with the AMBER99sb+bsc0 force field and previously developed CUFIX corrections for ion–ion^42^, ion–protein^42^, ion–DNA^42^, and protein–protein^50^ interactions to build a new force field set, AMBER99sb+bsc0+cufix. The complete force field set is available online at http://bionano.physics.illinois.edu/CUFIX.

### Diffusion of PCNA on DNA using the optimized force field set

Using AMBER99sb+bsc0+cufix, we produced a 2 μs-long trajectory of PCNA on DNA. Unlike the simulations using the standard force fields, PCNA showed dynamic movements both translationally and rotationally (Fig. 2C,E and Supplementary Movie 4). During 2 μs, the net translation of PCNA was about 5 nm (Fig. 2C), resulting in *D*_t_ = 3.4 nm^2^/μs that quantitatively agrees with the experimental *D*_t_ = 2.4 nm^2^/μs within the experimental standard deviation (Fig. 2D). This quantitative agreement suggests that the newly developed corrections dramatically improved the realism of MD simulations. Although the instantaneous tilt of PCNA rapidly fluctuated between 5° and 25° during 2 μs, the average tilt 13° was close to 15° observed in the crystal structure^19^ (Fig. 2F). Contacts between basic K/R residues and DNA phosphates were dynamic and stochastic, with the lifetime mostly shorter than 10 ns (Fig. 3D). The average K/R–DNA contact number was consistent with the contact number 7 observed in the crystal structure^19^ (Fig. 2G).

## Conclusion

Here, we showed that the standard AMBER and CHARMM force fields overestimate the attractive interactions between basic K/R residues and DNA phosphate groups, overstabilizing the contacts at the PCNA–DNA interfaces. To balance those charge–charge interactions, we optimized the LJ parameters for the guanidinium nitrogen–phosphate oxygen and amine nitrogen–phosphate oxygen atoms pairs using the experimental osmotic pressure data of model compounds. By using the optimized LJ parameters, we were able to match the diffusion coefficient of PCNA quantitatively. Given that DNA-binding proteins generally utilize abundant K/R residues to stabilize the protein–DNA contacts, we expect that our optimized parameters will have a broad impact on the simulations of protein–DNA complexes.

Although the main scope of this manuscript is the validation of the K/R–DNA interactions, it is worthwhile to mention the diffusion mechanism of PCNA briefly. Through the single-molecule tracking experiments, Kochaniak et al. proposed the translational and helical modes, in which PCNA undergoes a fast sliding motion with low translation-rotation coupling and a slow sliding motion following the helical path of DNA backbone, respectively^22^. Based on the diffusion coefficient close to the experimental upper limit (Fig. 2D) and the low translation-rotation coupling (compare Fig. 2C and 2E), it seems that our simulation using AMBER99sb+bsc0+cufix is consistent with the translational mode. Because the ideally linear DNA used in our simulations may prefer the fast translation mode to the slow rotational mode, simulations on realistically curved DNA might be necessary for the future study. Recently, based on the MD simulations using the standard AMBER force field, De March et al. proposed the cogwheel mode, in which PCNA slides along the helical path of DNA by tracking the DNA backbone with a fixed 30° tilt of PCNA maintained by the PCNA–DNA contacts^19^. Based on our validation of the standard AMBER force fields and the dynamically changing tilt and PCNA–DNA contacts using AMBER99sb+bsc0+cufix, it seems that our MD simulation does not support the cogwheel mode.

## Supporting information

Supplementary Movie 1. MD trajectory using the standard AMBER99sb+bsc0 force field (Movie)

Supplementary Movie 2. MD trajectory using the standard AMBER14sb+bsc1 force field (Movie)

Supplementary Movie 3. MD trajectory using the standard CHARMM36m force field (Movie)

Supplementary Movie 4. MD trajectory using the AMBER99sb+bsc0+cufix force field (Movie)

## ASSOCIATED CONTENT

### Supporting Information

The following files are available free of charge.

Supplementary Movie 1. MD trajectory using the standard AMBER99sb+bsc0 force field (Movie)

Supplementary Movie 2. MD trajectory using the standard AMBER14sb+bsc1 force field (Movie)

Supplementary Movie 3. MD trajectory using the standard CHARMM36m force field (Movie)

Supplementary Movie 4. MD trajectory using the AMBER99sb+bsc0+cufix force field (Movie)

## Notes

The authors declare no competing financial interest.

## ACKNOWLEDGMENT

J.Y. gladly acknowledge computer time provided through the HQ cluster at the Institute for Basic Science.

